# AxSpA joint tissue is characterised by HLA-DR^+^ tissue resident memory (TRM) and killer cell immunoglobulin-like receptor (KIR)^+^ CD8^+^ T cell subsets

**DOI:** 10.1101/2024.10.09.617402

**Authors:** Feng Liu, Hui Shi, Jiewei Chen, Ben Kendrick, Dajiang Du, Paul Bowness, Qiang Tong, Liye Chen

**Affiliations:** Botnar Research Centre, Nuffield Department of Orthopaedics, Rheumatology and Musculoskeletal Sciences, University of Oxford, Oxford, UK; Department of Rheumatology and Immunology, Shanghai Sixth People’s Hospital Affiliated to Shanghai Jiao Tong University School of Medicine, Shanghai, China; Shanghai Sixth People’s Hospital Affiliated to Shanghai Jiao Tong University School of Medicine, China; Nuffield Orthopaedic Centre, Oxford University Hospitals Trust, Oxford, United Kingdom; Nuffield Department of Orthopaedics, Rheumatology and Musculo-skeletal Science, University of Oxford, Oxford, United Kingdom

## Abstract

**Objective:** Axial Spondyloarthritis (AxSpA) is a common inflammatory arthritis with *HLA-B*27* as the major genetic risk. Recent discoveries of AxSpA-specific T cell receptor (TCR) motifs and the self and bacterial peptides that they recognize support a pathogenic role of CD8^+^ T cells. Despite of previous work on synovial fluid, the characteristics of CD8 cells in joint tissue are currently unknown.

**Method:** Synovial tissues from 5 AxSpA patients were used for single cell RNA sequencing (scRNA-seq). Paired TCR sequencing was carried out for 2. The abundance of KIR^+^CD8^+^ T cells in the blood from 9 AxSpA patients and 10 healthy controls was measured using flow cytometry. The expression of naïve and memory T cell markers (CCR7, CD45RA and CD45RO) were compared between KIR^+^ and KIR^-^ CD8 cells.

**Results:** We observed conventional, *TRAV1-2*^*+*^ mucosal-associated invariant T (MAIT) cell and *MKI67*^*+*^ proliferating cell populations in synovium. Following sub-clustering of conventional CD8^+^ T cells, *HLA-DR*^*+*^ tissue resident memory (TRM), circulating, *KIR*^*+*^ and *FCGR3A*^*+*^ (encoding CD16) cell subsets were observed. *HLA-DR*^*+*^ TRM and *KIR*^*+*^ cells were clonally expanded and exhibited distinct transcriptional features, enriched for T cell activation pathways and natural killer (NK) cell-mediated cytotoxicity pathway respectively. Lastly, KIR^+^CD8^+^ T cells were increased in AxSpA blood and enriched for CD45RA^+^CCR7^-^ T_EMRA_ cells.

**Conclusion:** Here we present the very first transcriptomic profiling of CD8^+^ T cells in synovium tissue and highlight potential roles of *HLA-DR*^*+*^ TRM and *KIR*^*+*^ cells in AxSpA pathology. This study adds novel insights to the disease mechanisms and offers new therapeutic opportunities.

## INTRODUCTION

Axial Spondyloarthritis (AxSpA) is a common immune-mediated inflammatory arthritis with *HLA-B*27* as the major genetic risk. A critical pathogenic role for HLA-B*27-restricted CD8^+^ T cells in AxSpA has long been mooted. This has recently received further support with the discovery of T cell receptor (TCR) motifs that are enriched in AxSpA synovial fluid (SF) (1)(2). Notably, TCRs with these motifs have been found to recognize both self and bacterial antigenic peptides presented by HLA-B*27 (3). In line with the frequent incorporation of the β chain TRBV9 by these TCR motifs, a case study has shown the potential clinical efficacy of an antibody depleting TRBV9^+^ T cells (4).

CD8^+^ tissue resident memory (TRM) T cells have been implicated in host defence, cancer and autoimmune diseases. CD8^+^ T cells with TRM features have been found in synovial fluid (SF) and gut tissue from AxSpA patients and the inflamed spine of monkeys with AxSpA-like disease (2,5–7). Additionally, CD8^+^ T cells expressing inhibitory killer cell immunoglobulin-like receptors (KIRs) are expanded in the blood and inflamed tissues of patients with a variety of autoimmune diseases and capable of eliminating pathogenic gliadin-specific CD4^+^ T cells from the leukocytes of celiac disease patients in vitro (8). Considering the ability of HLA-B*27 heterotrimers and homodimers to interact with KIR3DL1 and KIR3DL2 respectively (9), it is intriguing to ask if such KIR^+^CD8^+^ T cells have a role in disease in AxSpA.

Here we use single-cell RNA/TCR-sequencing (scRNA/TCR-seq) to study CD8^+^ T cells from AxSpA synovial tissue and observe the clonal expansion of two subsets: HLA-DR^+^ TRM and KIR^+^ cells. Further, we show that the frequency of KIR^+^CD8^+^ T cells are increased in AxSpA blood and are enriched for the terminally differentiated CD45RA^+^CCR7^-^ effector memory cells (T_EMRA_).

## MATERIALS AND METHODS

### Patient Recruitment

Fresh synovial tissue was obtained at arthroplasty was collected from patients attending the Oxford University Hospitals (OUH) or Shanghai Sixth People’s Hospital. Peripheral blood was collected from patients attending the Oxford University Hospitals National Health Service Foundation Trust (OUH). Both tissue and blood samples were obtained with appropriate consent and ethical approval. All patients included in this study met the criteria of the Assessment of Spondyloarthritis International Society (ASAS) for AxSpA. The demographics of AxSpA patients recruited for this study are shown in Table S1.

### scRNA-seq experiments and data analysis

We obtained synovial tissues from joint replacement surgeries, which were subsequently washed, and the fat pads were removed. Single-cell suspensions were then generated from the tissue using Liberase digestion. The isolated cells were either directly profiled using the Chromium Single Cell Gene Expression assay (10X Genomics) or sorted into live CD3+ and CD3-populations using Fluorescence-activated Cell Sorting (FACS). When FACS was employed, the CD3+ and CD3-populations were loaded into separate channels of the Chromium chip, where 5’ gene expression and immune repertoire profiling assays were conducted for the simultaneous detection of TCR and transcriptomics. DNA libraries generated from the 10X Chromium assay were sequenced with a depth of over 50,000 reads per cell on a NovaSeq 6000.

The obtained reads were mapped to the hg38 reference genome to generate gene expression matrices using CellRanger. The expression matrix was then analyzed with the Seurat R package. Initially, low-quality cells were removed based on criteria such as the number of detected genes, the percentage of mitochondrial RNA among total UMIs, and the total number of UMIs. Subsequent steps, including data normalization, scaling, integration using the CCA algorithm, dimension reduction, and differential gene expression analysis, were performed following the Seurat vignette. Unless otherwise specified, default parameters were used for each function.

### Flow cytometry phenotyping of KIR^+^CD8^+^ T cells

Isolated PBMCs from AxSpA patients or healthy controls were stained with the following antibodies: CD3-BV786 (clone OKT3; BioLegend), CD4-BV605 (clone RPA-T4; BioLegend), CD8-BV711 (clone RPA-T8; BioLegend), CD45RA-BV650 (clone HI100; BioLegend), CCR7-PE-Cy7 (clone G043H7; BioLegend) and pan-KIR-PE. The pan-KIR-PE was constituted of five antibodies against KIR2DL1 (clone FAB1844P; R&D), KIRDL2-3 (clone DX27; BioLegend), KIR3DL1 (clone DX9; BioLegend), KIR3DL2 (clone FAB2878A; R&D) and KIR2DL5 (clone UP-R1; BioLegend). To exclude dead cells, samples were stained with LIVE/DEAD™ Fixable Violet Dead Cell Stain Kit (L34955, Invitrogen).

### Patient and Public Involvement

We will actively engage the patients with AxSpA through the Oxford Patient Engagement Network for Arthritis and Musculoskeletal Conditions (OPEN ARMS, https://www.ndorms.ox.ac.uk/get-involved/open-arms-1/open-arms) and the Botnar AxSpA day. We will discuss with patients about the impact of findings from this study and co-develop a dissemination plan to maximize the potential benefits of our findings to patients.

### Statistics

The level of statistical significance was assessed by unpaired test (Figure 3B) or paired Wilcoxon test (Figure 3D). The differences were considered statistically significant at P<0.05. All statical analyses and summarized graphs were performed using GraphPad Prism 10.3.0.

### Ethics approval

Venous blood was obtained under protocols approved by the Oxford Research Ethics committee (ethics reference number 06/Q1606/139). Synovial tissue was obtained under protocols approved by the Oxford Research Ethics committee (ethics reference number 06/Q1606/139) and Ethics Committee of Shanghai Sixth People’s Hospital (2024-KY-132).

### Data availability

Data are available upon reasonable request.

## RESULTS

### Four subsets of conventional CD8^+^ T cells are found in AxSpA synovial tissue

We studied synovium from 5 AxSpA patients undergoing arthroplasty (four hip and one knee, Table S1). A total of 5663 CD8^+^ T cells were categorized into four clusters, two conventional CD8+ T cell clusters, a *TRAV1-2*^+^ mucosal-associated invariant T (MAIT) cell cluster and a *MKI67*^+^ proliferating cluster (Figure 1A and B, Figure S1). Conventional CD8^+^ T cells were extracted and sub-clustered into four subsets (Figure 1C). The largest subset was characterized by expression of *HLA-DRA* and of the TRM marker (*CXCR6)* together with low levels of the circulating markers (*CCR7* and *S1PR1*) (Figure 1D). In contrast the “circulating” cluster had the opposite expressional pattern of these genes. The third and fourth clusters featured expression of *KIR* (*KIR2DL3* and *KIR3DL2*) and *FCGR3A* (encoding CD16) respectively. These four populations were present in very similar proportions in each of the 5 AxSpA patients (Figure 1E).

**Figure 1.**
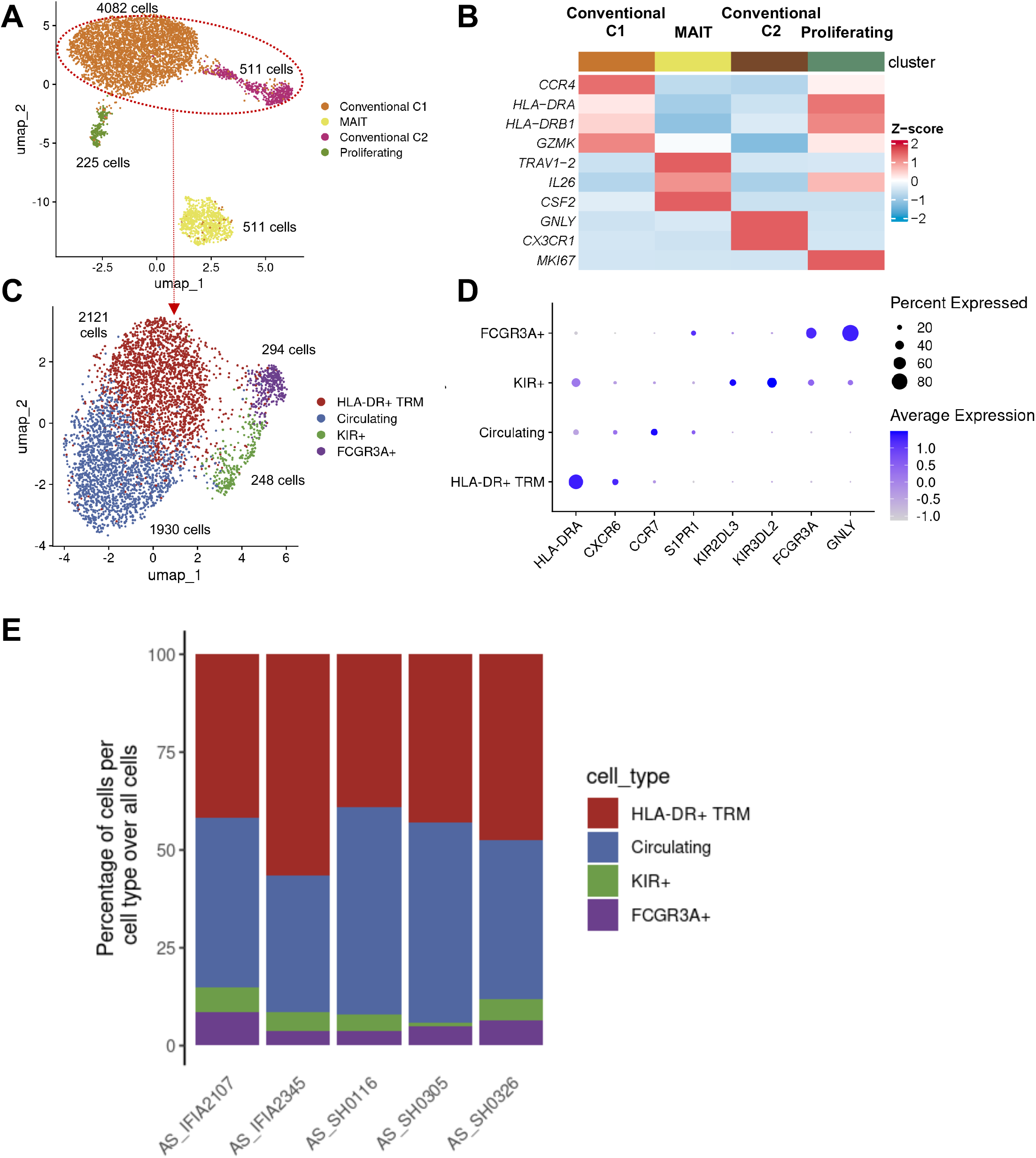
Single-cell transcriptomic characterization of CD8+ T cells in AxSpA synovial tissue. (A) UMAP visualization of transcriptionally distinct populations CD8+ T cells from synovial tissues of 5 AxSpA patients (4 hip, 1 knee). (B) Heatmap of normalized and scaled expression of marker genes of different cell subsets. (C) UMAP visualization of subsets of conventional CD8+ T cells. (D) Average expression of marker genes by different conventional CD8+ T cell subsets. Dot size represents the percentage of cells with detected expression of corresponding genes. (E) Percentage of four subsets in each patient is shown. Synovium was collected from knee (AS_IFIA2345) or hip (rest four samples).

### The HLA-DR^+^ TRM and KIR^+^ cell subsets are clonally expanded and exhibit distinct transcriptional features

We then asked if clonal T cell expansion is present in any of these subsets. To this end, we carried out paired RNA and TCR sequencing for two AxSpA patients and observed the enrichment of top TCR clonotypes within the *HLA-DR*^+^ TRM and *KIR*^+^ clusters (Figure 2A). This finding drove us to define the transcriptional features of these two clusters. Figure 2B shows the upregulation of genes related to activation (*HLA class II*), cytotoxicity (*GZMH, GZMA* and *PFN1*) and tissue residency (*CXCR6* and *TOX2*) in the *HLA-DR*^+^ TRM. In contrast, The *KIR*^*+*^ cluster upregulated multiple inhibitory KIRs together with NK-related genes (*KLRC2, KLRC3* and *KLRD1*) and *IKZF2* (encoding transcription factor Helios and previously reported in KIR^+^ CD8 (8)) (Figure 2C). Pathways analysis using top feature genes (log2 fold change >1) further suggested that these two subsets are transcriptional distinct (Figure 2 D and E). In addition to the expected MHC class II pathway, the *HLA-DR*^+^ TRM cell population was also linked to multiple pathways related to T cell activation. The *KIR*^*+*^ cluster was strongly associated with natural killer (NK) cell-mediated cytotoxicity pathway.

**Figure 2.**
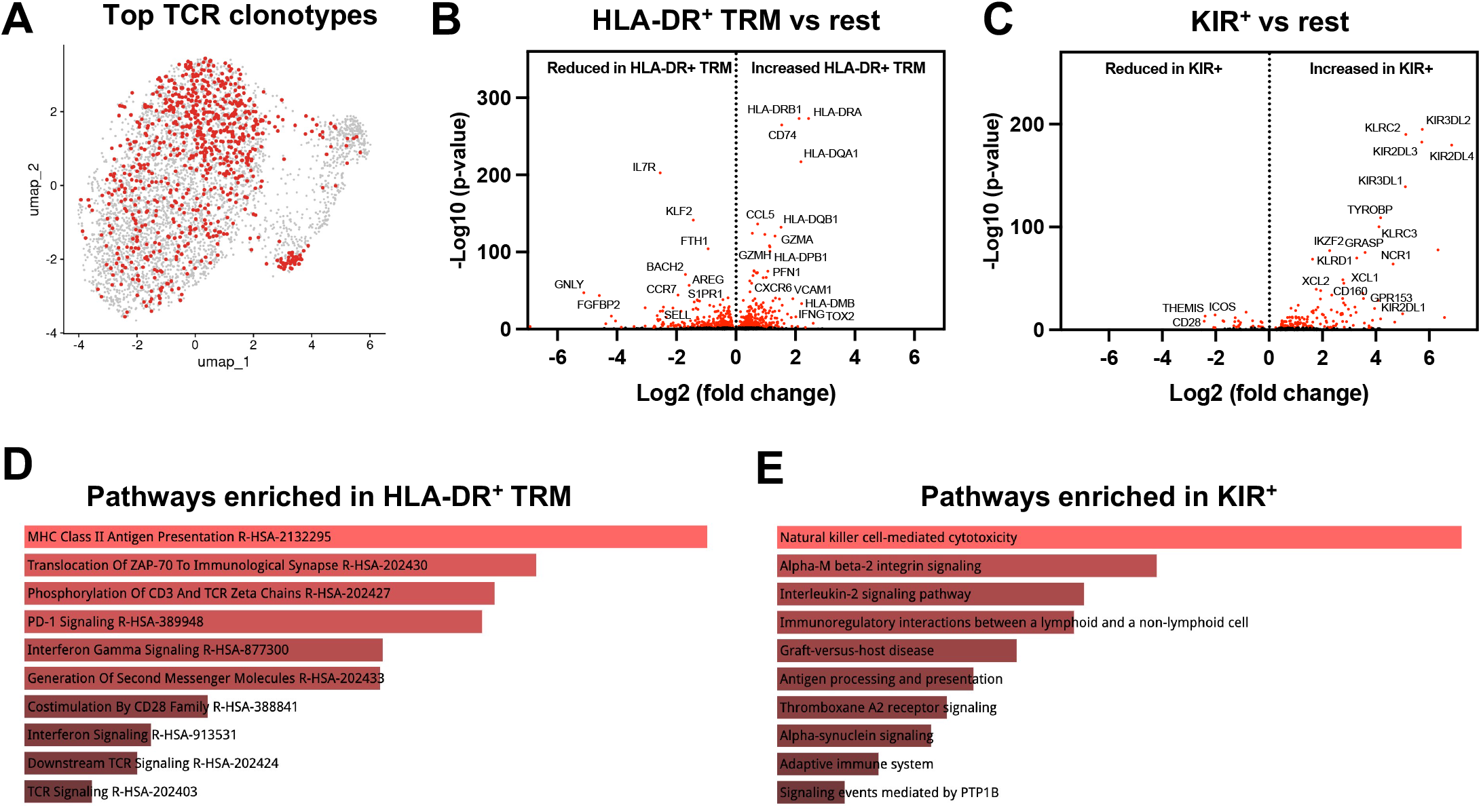
Phenotypic characterization of HLA-DR+CD8+ TRM and KIR+CD8+ T cells in AxSpA synovial tissue. (A) the top 6 most frequent TCR clonotypes from 2 AxSpA patients (red dots) projected onto overall CD8+ T cell UMAP. (B and C) Volcano plots showing genes differentially expressed in HLA-DR+ TRM (B) and KIR+ cells (C) compared to other cells. Pathway enrichment analysis (Reactome and BioPlanet) carried out using top feature genes (Log 2 fold change >1) are shown for HLA-DR+ TRM (D) and KIR+ cells (E) respectively. Only significantly enriched pathways are shown (p< 0.05).

### KIR^+^CD8^+^ T cells are increased in AxSpA blood and enriched for CD45RA^+^CCR7^-^ T_EMRA_

Elevated frequencies of blood KIR^+^CD8^+^ T cells have been shown in systemic lupus erythematosus, multiple sclerosis and celiac disease (8). This drove us to ask if KIR^+^CD8^+^ T cells are expanded in AxSpA blood. Figure 3A shows that KIR^+^ cells are increased in AxSpA CD8^+^ T cells compared to healthy controls (Figure 3 A and B). We then carried out further immune phenotyping using makers for naïve and memory T cells and found these KIR^+^ AxSpA CD8^+^ T cells were predominantly CCR7-CD45RA+ indicating a T_EMRA_ phenotype (Figure 3C and D).

**Figure 3.**
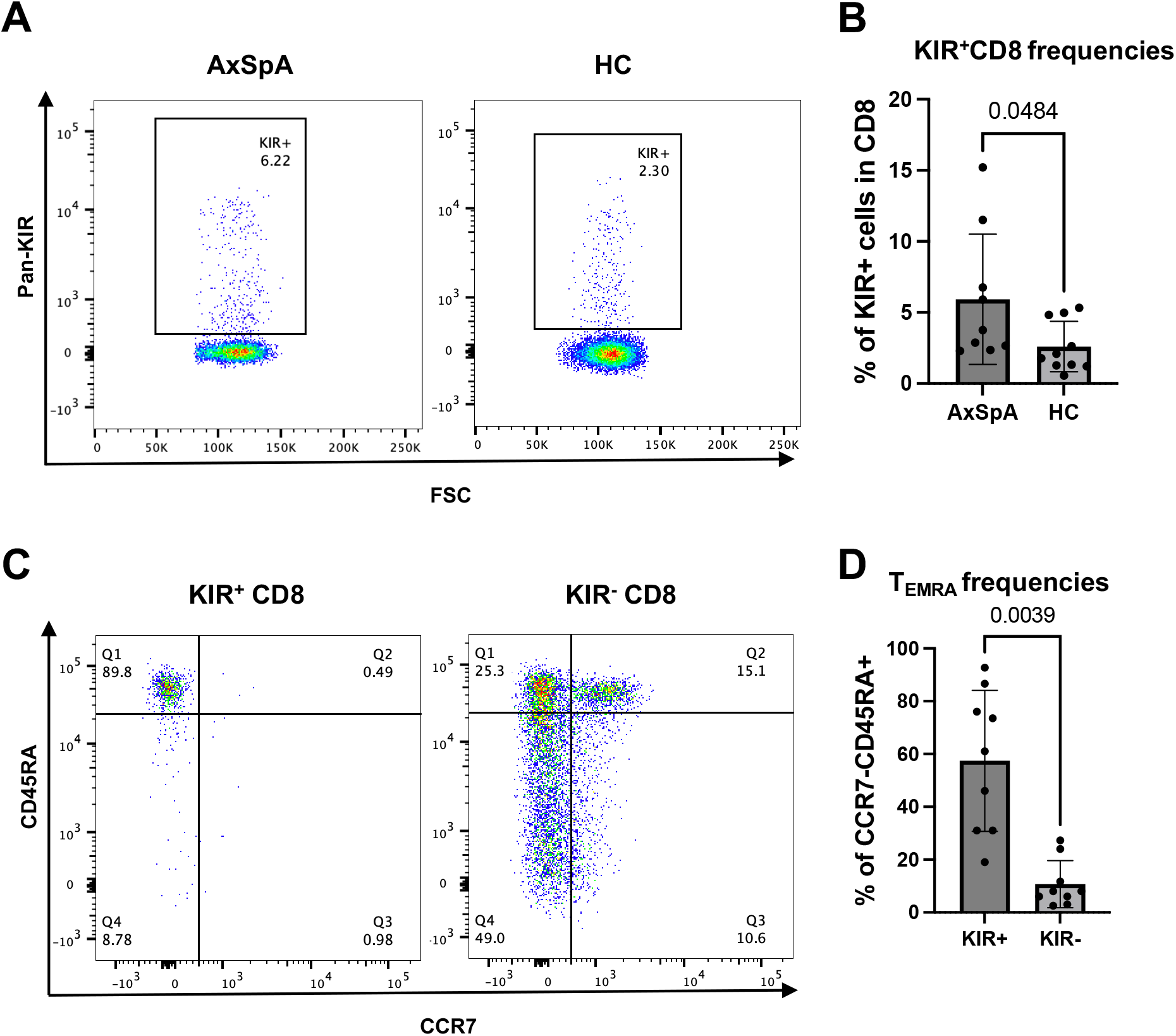
Phenotypic characterization of HLA-DR+CD8+ TRM and KIR+CD8+ T cells in AxSpA synovial tissue. (A) Staining and comparison of KIR+ CD8+ T cells between AxSpA (n=9) and HC (n=10). Summary graph is shown in (B). (C) Staining and comparison of CCR7-CD45RA+ TEMRA cells between KIR+ and KIR-CD8 in AxSpA blood (n=9). Summary graph is shown in (D). Unpaired test and paired Wilcoxon test used for B and D respectively. p < 0.05 considered significant.

## DISCUSSION

In this study, we transcriptionally profile the synovial tissue CD8^+^ T cells at the single cell level for the first time. The previously reported CD8^+^ TRM in synovial fluid is also found in synovial tissue. Notably we also find for the first time the presence of *KIR*^*+*^ *CD8*^*+*^ T cell, a recently discovered novel type of regulatory T cells (8), in the synovium. Interestingly, the flow cytometry phenotyping of AxSpA blood cells reveal an increased frequency of KIR^+^ CD8^+^ T cells, which strongly enrich the CD45RA+CCR7-T_EMRA_ cells.

Whether and how KIR^+^CD8^+^ T cells contribute to AxSpA pathology are not answered by our study. These cells have been shown to specifically eliminate auto-reactive CD4^+^ T cells in several autoimmune diseases through their cytotoxicity function (8). With interleukin 17 (IL-17) being a key driver of AxSpA inflammation, it would be interesting to test if KIR^+^CD8^+^ T cells kill CD4^+^ Th17 cells in AxSpA.

How KIR^+^CD8^+^ T cells recognize auto-reactive CD4^+^ T cells is not completely understood. However, antibody blocking of either classical HLA class molecules or HLA-E have been shown to compromise this regulatory function in the in vitro assay (8). Considering the previous data showing the binding of HLA-B*27 with KIR3DL1 and KIR3DL2 (9), it is plausible to hypothesize that HLA-B*27 could affect the function and generation/maintenance of KIR^+^CD8^+^ T cells.

The limitations of this study include: 1) Due to funding and sample accessibility challenges, we have not been able to sequence tissue from control donors (such as patients with Osteoarthritis). 2) Due to the size of tissue sample that we acquired, we have not been able to obtain enough number of HLA-DR^+^ TRM and KIR^+^ CD8 cells from joint for functional studies.

In summary, our data confirm the presence in AxSpA synovial tissue of CD8^+^ T cells with an activated TRM phenotype previously described in AxspA synovial fluid, and show for the first time the presence of a *KIR*^+^*CD8*^+^ T cell population. KIR^+^CD8^+^ T cells are also increased in AxSpA blood supporting a potential role of in AxSpA. Our findings suggest that more than one CD8^+^ T cell population may be involved in AxSpA pathogenesis and provide new opportunities for therapeutic interventions.

## Author contributions

FL, PB, QT and LC designing research studies, FL, HS, JC, BK and DD conducting experiments, FL and HS acquiring data, FL, HS and LC analyzing data, FL, PB, QT and LC writing the manuscript.

## Acknowledgements

This work was funded by a Versus Arthritis career development award to LC 22053. P.B. is funded by the National Institute for Health Research (NIHR) Oxford Biomedical Research Centre (BRC). The views expressed are those of the author(s) and not necessarily those of the NHS, the NIHR or the Department of Health.

## FIGURE LEDENDS

**Supplementary Table 1.** Demographics of AxSpA patients recruited.

| **Supplementary Table 1. Demographics of AxSpA patients recruited** |  |  |
| --- | --- | --- |
|  | **Tissue samples (n=5)** | **Blood samples (n=9)** |
| Age, mean (range) years | 61.8 (50-79) | 37.9 (19-58) |
| Sex, male/female | 4/1 | 5/4 |
| HLA-B27, +/-/unknown | 4/0/1 | 7/1/0 |
| Sacroiliitis by X-Ray (Yes/No) | 5/0 | NA |
| CRP, mean (range) | 22.38 (0.16-38.37) | 13.56 (0.5-51.7) |
| BASDAI, mean (range) | 3.36 (1.6-7.2) | 4.76 (1.9-7.24) |
| NASID current (Yes/No) | 4/1 | 6/3 |
| Biologic DMARD therapy current (Yes/No) | 1/4 | 2/7 |
| CRP: C-reactive protein; BASDAI: Bath Ankylosing Spondylitis Disease Activity Index; NSAIDs: Non-steroidal anti-inflammatory drugs; DMARD: Disease-modifying antirheumatic drugs. |  |  |

**Figure S1.**
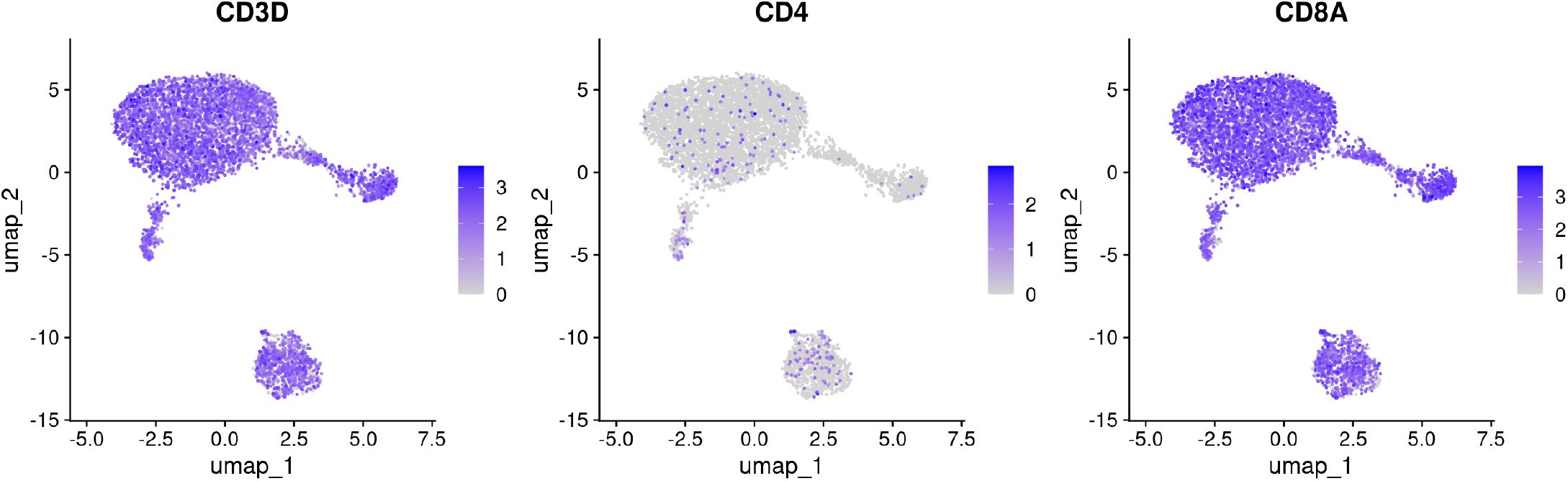
Characterization of AxSpA synovial tissue CD8+T cells. UMAP visualization of normalized expression of *CD3D, CD4* and *CD8A* of 5663 T cells from 5 AxSpA tissues (4 hip, 1 knee).

**Figure S2.**
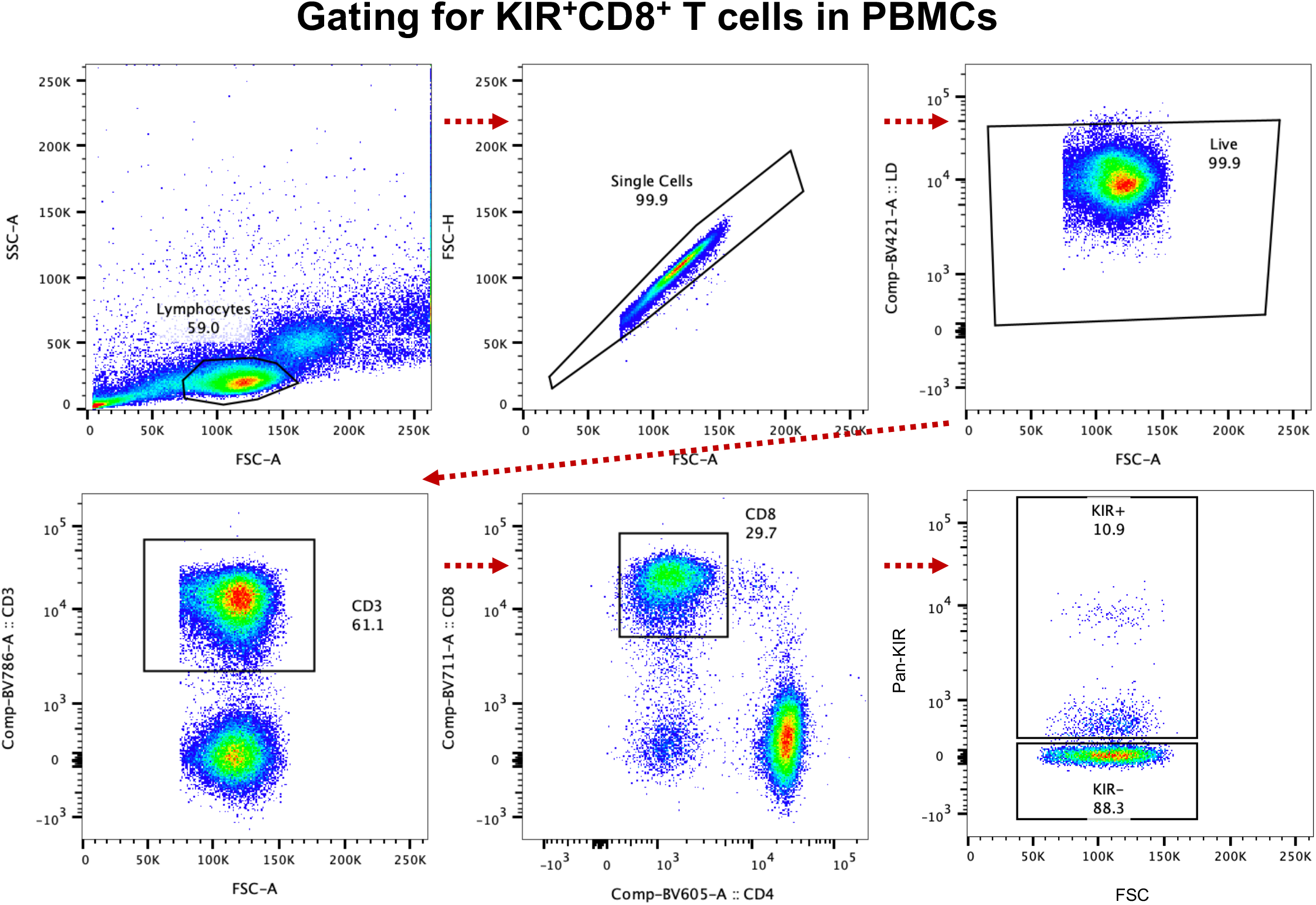
FACS gating strategy for phenotyping KIR^+^CD8^+^ T cells from AxSpA and healthy PBMCs. Staining of PBMCs from a AxSpA patient is shown.

## REFERENCES

1. Faham M, Carlton V, Moorhead M, Zheng J, Klinger M, Pepin F, et al. Discovery of T Cell Receptor β Motifs Specific to HLA–B27–Positive Ankylosing Spondylitis by Deep Repertoire Sequence Analysis. Arthritis Rheumatol 2017;69:774–784.

2. Deschler K, Rademacher J, Lacher SM, Huth A, Utzt M, Krebs S, et al. Antigen-specific immune reactions by expanded CD8+ T cell clones from HLA-B*27-positive patients with spondyloarthritis. J Autoimmun 2022;133:102901.

3. Yang X, Garner LI, Zvyagin IV, Paley MA, Komech EA, Jude KM, et al. Autoimmunity-associated T cell receptors recognize HLA-B*27-bound peptides. Nature 2022:1–7.

4. Britanova OV, Lupyr KR, Staroverov DB, Shagina IA, Aleksandrov AA, Ustyugov YY, et al. Targeted depletion of TRBV9+ T cells as immunotherapy in a patient with ankylosing spondylitis. Nat Med 2023;29:2731–2736.

5. Guan T, Bian Z, Gao H, He Y, Yuan J, Wan H, et al. Altered CD8+ T cell subpopulation in the bone marrow microenvironment of cynomolgus monkeys with spontaneous ankylosing spondylitis. Ann Rheum Dis 2024:ard-2024-226018.

6. Guggino G, Rizzo A, Mauro D, Macaluso F, Ciccia F. Gut-derived CD8+ tissue-resident memory T cells are expanded in the peripheral blood and synovia of SpA patients. Ann Rheum Dis 2021;80:e174–e174.

7. Qaiyum Z, Gracey E, Yao Y, Inman RD. Integrin and transcriptomic profiles identify a distinctive synovial CD8+ T cell subpopulation in spondyloarthritis. Ann Rheum Dis 2019;78:1566.

8. Li J, Zaslavsky M, Su Y, Guo J, Sikora MJ, Unen V van, et al. KIR+CD8+ T cells suppress pathogenic T cells and are active in autoimmune diseases and COVID-19. Science 2022:eabi9591.

9. Kollnberger S, Bird L, Sun M-Y, Retiere C, Braud VM, McMichael A, et al. Cell-surface expression and immune receptor recognition of HLA-B27 homodimers. Arthritis Rheumatism 2002;46:2972–2982.

